# Insights into Biomarker Correlations in Relation to Stress: A Preliminary Analysis

**DOI:** 10.1101/2023.09.16.557862

**Authors:** Annika Lundström, Caterina Mandalios

**Affiliations:** University of Helsinki

## Abstract

The objective of this study was to analyze specific biomarkers of interest and find correlations between them using hormonal and cardiovascular measurements. The aim was to explore these physiological patterns during a typical weekday. While this research provides valuable insights into physiological patterns and their potential impact on mental performance, it’s essential to note that the study did not aim to diagnose, treat, or propose specific medical interventions based on the findings. Instead, the focus was solely on observational data collection and analysis for the purpose of exploring correlations and understanding daily physiological fluctuations in the context of stress and cognitive performance. Furthermore, the team observed how a low-level stress-triggered cognitive task could alter, and possibly impair, mental performance due to fluctuations in these selected data points. Five (5) adult, healthy participants (aged between 20 and 35 years) volunteered for the small-scaled study (two male and three female).

In the current study, participants with larger hormonal and cardiovascular fluctuations during a 5-minute stress-induced task did not achieve as well as those with more stable physiological measurements. Interestingly, a significant peak in cortisol levels was detected during the mental task, likely due to the stress-induced environment.

The current, and vastly limited methods used for at-home, hormonal measurements, as well as the slightly unreliable sensor technology used in general pulse oximeters measuring blood oxygen saturation in this trial, indicate the great need to develop new solutions. This includes innovations that would make health monitoring more convenient, while at the same time emphasizing improved mental well-being in daily situations, all without having to continuously wear a gadget to obtain precise insights about multiple health metrics.

## Introduction

Research has consistently indicated correlations between hormonal fluctuations, cardiovascular rate changes, and cognitive decline [1], [2]. Understanding the potential mechanisms underlying this association is essential, in order to get a comprehensive grasp of cognitive health. With the rapid advancement of digital technologies in healthcare, this has opened new possibilities in real-time health monitoring and personalized interventions [3]. Central to these developments is the vastly growing field of Health Technology (*HealthTech*), where devices integrated with innovative technologies are transforming the delivery and convenience of healthcare [3]. Detecting early indications of prolonged, heightened stress levels and other physiological imbalances, could ease and prevent a vast majority of people suffering from long-term health complications such as depression, anxiety and other mental health-related issues, metabolic disorders, and compromised immune function [4],[5], thereby improving overall public health and reducing medical intervention costs.

Cortisol, a glucocorticoid hormone, is synthesized and secreted by the adrenal cortex in response to physiological or psychological stressors [6]. Alongside epinephrine (adrenaline), it plays a pivotal role in the body’s acute stress response [6]. During short-term stress exposure, increased cortisol levels can have adaptive benefits by enhancing physical and cognitive performance and modulating inflammatory responses [7]. However, chronic exposure to elevated cortisol concentrations has shown to have damaging effects on various physiological systems, for example, hyperglycemia, enhanced risk of Type II Diabetes, or cause a prolonged elevation in blood pressure, thereby increasing the risk of cardiovascular diseases and cognitive decline [8],[9]. Early detection of abnormalities in cortisol levels, usually requires a comprehensive physical examination and blood sampling taken by a physician at a clinical institution. Therefore, the necessity of making alternative diagnostic methods to become more easily accessible is highly of interest.

Alongside cortisol, other important hormonal imbalances of e.g., progesterone, estrogen, and testosterone, can often go unnoticed and untreated for a long time. Dysregulations in these steroid hormones can similarly develop into chronic conditions, negatively affecting mental well-being, reproductive health, bone density, and cardiovascular function. [10], [11].

This study measured and analyzed cardiovascular and hormonal data from healthy, adult participants in their normal living conditions during a typical weekday. The aim was to find possible correlations between these biomarkers and to develop a more profound understanding of how stress may interfere with mental performance and overall behavior patterns.

By conducting an initial research trial for the purpose of validating the ideas and theories collected prior to the study regarding the brain’s neuroplastic abilities and cognitive adaptation processes, more concrete results of these aspects were successfully gained. Supported by previous research within the neuroscientific field, this study created a good foundation for the team’s further advancements within health technology.

The findings from this study, to be discussed in subsequent sections, provide valuable insights into how rapid fluctuations in cortisol, blood oxygen saturation, and resting heart rate influence one’s state of mind during a stress-induced task, as well as the importance of raising the general awareness of potential imbalances in steroid hormone levels. Additionally, this research trial highlights the significance of developing effective and concise methods for stress management in order to optimize one’s own daily tasks and routine.

This article further discusses the methodology adopted, analysis performed, results obtained, limitations of the study and the implications on the future development of a HealthTech device for cognitive enhancement.

## Materials and Methods

### Subjects

Five (5), healthy adults volunteered for the initial proof-of-concept study, aged between 20 and 35 years. Prior to the study, written informed consent was obtained from the individuals and the participants were briefed about the study’s objectives and their right to withdraw from the study at any point without any consequences. Confidentiality of their personal details and health information was assured according to the GDPR requirements, and any queries they had were addressed before commencing the study. The participants’ lifestyles were similar in the sense that everyone was working either remotely or in an office space at their full-time jobs during the time of the study, which made the comparison in biomarker results more consistent and reduced potential confounding variables related to differences in daily activity or situational stressors.

### Rationale for the Selected Measurements

Supported by previous research made in neuroscience, the brain and the body have been shown to be closely interconnected [12], and there are many physiological factors that particularly regulate how well the brain can receive and process information in different situations throughout the day [13].

By analyzing how certain hormonal and cardiovascular rates interact with each other, one of the aims of the trial was to gain useful insights about how the body’s inner rhythm fluctuates, especially when exposed to a considerable stressful environment. The results will serve as a reference for the research team’s future endeavors.

### General Protocol

The participants of the study were instructed to provide simple saliva samples in collection tubes at different times during one, normal weekday in their lives. Additionally, they were informed to wear a smart ring the night before as well as during the day of measurements to detect cardiovascular rates.

Most of the salivary cortisol samples were collected early in the morning (upon waking, 30 minutes after waking, and 1 hour after waking), with additional samples taken at various times throughout the day (see Figure 2). The steroid hormones, estrogen, and progesterone for female participants and respectively testosterone for male participants, were taken once upon wakening. The specific hormonal measurements included in this study are listed in Table 1.

**Table 1.** Summary of hormones analyzed in this study and the specific measurements taken for men and women, respectively.

|  | Cortisol | Testosterone | Estrogen | Progesterone |
| --- | --- | --- | --- | --- |
| male | x | x |  |  |
| female | x |  | x | x |

### Hormonal Collection

For the salivary measurements, the passive drool collection tubes from the at-home health test service, GetTested (www.gettested.fi) were used. Specifically for this study, cortisol levels were measured by all participants in the study. Additionally, either progesterone and estrogen (for women) or testosterone (for men) were analyzed. Supported by prior findings, these steroid hormones have been shown to have a distinct connection that can potentially interfere with cognitive function [14], [15]. Estrogen, progesterone, and testosterone are also considered as biomarkers which are not being tested on a regular basis due to the lack of accessibility.

### Laboratory analysis

After the hormonal collection was completed, the saliva sample kits used in this study were sent by the participants via mail to an ISO 13485-certified laboratory in Finland. For the analysis of the salivary samples, the laboratory facilities of the at-home health test service, GetTested, were used. The samples were analyzed using the Liquid Chromatography-Mass Spectrometry (LCMS) method. LCMS is a powerful technique that combines the separation capability of liquid chromatography with the quantitative and qualitative capabilities of mass spectrometry [16]. This method is particularly valuable for detecting and quantifying low concentration molecules in complex mixtures, like hormones in saliva [17].

Once the analysis was completed, the results were uploaded to the participants’ profiles on the GetTested’s online platform and reviewed together with the research team to be discussed and safely collected for further analysis.

### Cardiovascular Measurements

Cardiovascular data were collected from the participants using smart rings (the Oura Ring), which were worn during the previous night and throughout the duration of the hormonal measurements. For data privacy reasons, participants were required to already own an Oura ring to be eligible for the study. The metrics that were taken into consideration for this study were resting heart rate and Heart Rate Variability (HRV).

In addition to this, participants wore standard pulse oximeter devices on their pointer fingers before and after a 5-minute mental task. These devices were used to detect blood oxygen saturation levels and resting heart rate. The goal was to identify potential fluctuations in cardiovascular rates due to the stress-induced environment and, according to the team’s hypothesis, determine if these would have a potentially negative impact on mental performance [18]. The Pulse Oximeters used in this study are listed in Table 2.

**Table 2.** List of Pulse Oximeters Used in the Study and their product numbers.

|  |
| --- |
| Beurer PO35 Fingertip Pulse Oximeter Oxygen Saturation Monitor Product Nr. 454.31 |
| Beurer PO30 Fingertip Pulse Oximeter Oxygen Saturation Monitor, Product Nr. 44-5023 |
| Yonker YK-83C Fingertip Pulse Oximeter LOT: 201126011202 |

### Cognitive Assignment

As part of the research trial, a short, 5-minute, cognitive assignment was performed by all the participants in the early afternoon (between 12-2:00 pm, according to their time of waking up). Upon joint agreement with the participants, the task was executed prior to the specific cortisol measurement which was intended to be taken 5 hours after wakening.

As previously researched, cortisol levels should normally reach their peak during the first waking hours [19], and naturally decrease in the early afternoon moving into night [20]. The intention of timing the mental exercise in the middle of the day, was to potentially find a significant, abnormal spike in cortisol due to the stress-induced scenario the participants were placed in. The specific task was not disclosed in advance to maintain an element of surprise in the experiment. The assignment was done virtually through an online call via Microsoft Teams with the research team. However, it was harmless and primarily designed to challenge the participants’ mental concentration. The cognitive task itself consisted of three, time-limited exercises, in the form of mathematical problems, rapid country naming, and word scrabble challenges (listed in Table 4).

### Data Privacy, Encryption & GDPR Requirements

To maintain and prioritize participants’ privacy, the two, consecutive documents in this research study were screened and approved by an official lawyer in Helsinki, Finland. Due to the global, growing concerns about data leaks, adhering to the GDPR Requirements, and giving the consequent rights to the participants to forfeit the study at any point, are precautions that were taken carefully into consideration for the research study. To ensure the participants’ data remained intact during processing, analysis, and storage, a data encryption procedure was managed by one of the research group’s data analysts.

Before initiating the study, participants were informed that the collected research data might be used for further research within the same discipline or for research in other disciplines that support the findings. All these details were provided to the participants in the two previously mentioned documents, where both the study description and data safety measures were clearly outlined.

### Data Analysis

Data analysis and visualization of the research results were performed through utilizing the Bioinformatics Analysis Platform RNAcloudomics developed by the research team’s collaborative data analyst from the University of Helsinki (Das Roy, PhD in Biotechnology).

All figures presented in this study have been stored safely in an encrypted cloud server, only accessed by the research team in this study.

## Results

Despite the limited number of participants in the research trial, this pilot study offers initial insights and noteworthy observations into how stress, as reflected in cardiovascular and hormonal data, might influence cognitive performance. The participants who performed best in the cognitive assignments, had the most stable hormonal and cardiovascular levels prior to and throughout the research day, which is being visualized in the subsequent sections.

The study’s results align with earlier research [21], [22], [23] which indicates that individuals who handle stress well, whether consciously or not, are more likely to thrive in everyday situations.

### Salivary Hormonal Measurements

In this study, hormonal measurements were successfully collected and analyzed, with minor complications noted only for the estrogen-progesterone hormone result of one participant and a single cortisol measurement out of the 35 that were taken. This related to the fact that saliva is a sensitive biomarker [24], needing immediate analysis to preserve its inherent integrity and specificity. Consequently, it means that the at-home sampling kit’s results could have been affected by contamination, improper handling, or delays in delivery to the lab.

The testosterone results from the male subjects were both within the normal reference range as shown in Figure 1 (Reference values for men: 10 −230.9 pg/ml, as referenced from laboratory values and the website GetTested), but the differences between them were quite significant. There is some evidence to suggest that high levels of cortisol, especially over prolonged periods (chronic stress) [25] can lead to suppressed testosterone production [26]. However, since the two subjects of this study showed healthy results and the execution of this study being made on a limited, small scale, claims like these cannot be applied in this scenario without a more extensive examination being made.

**Figure 1.**
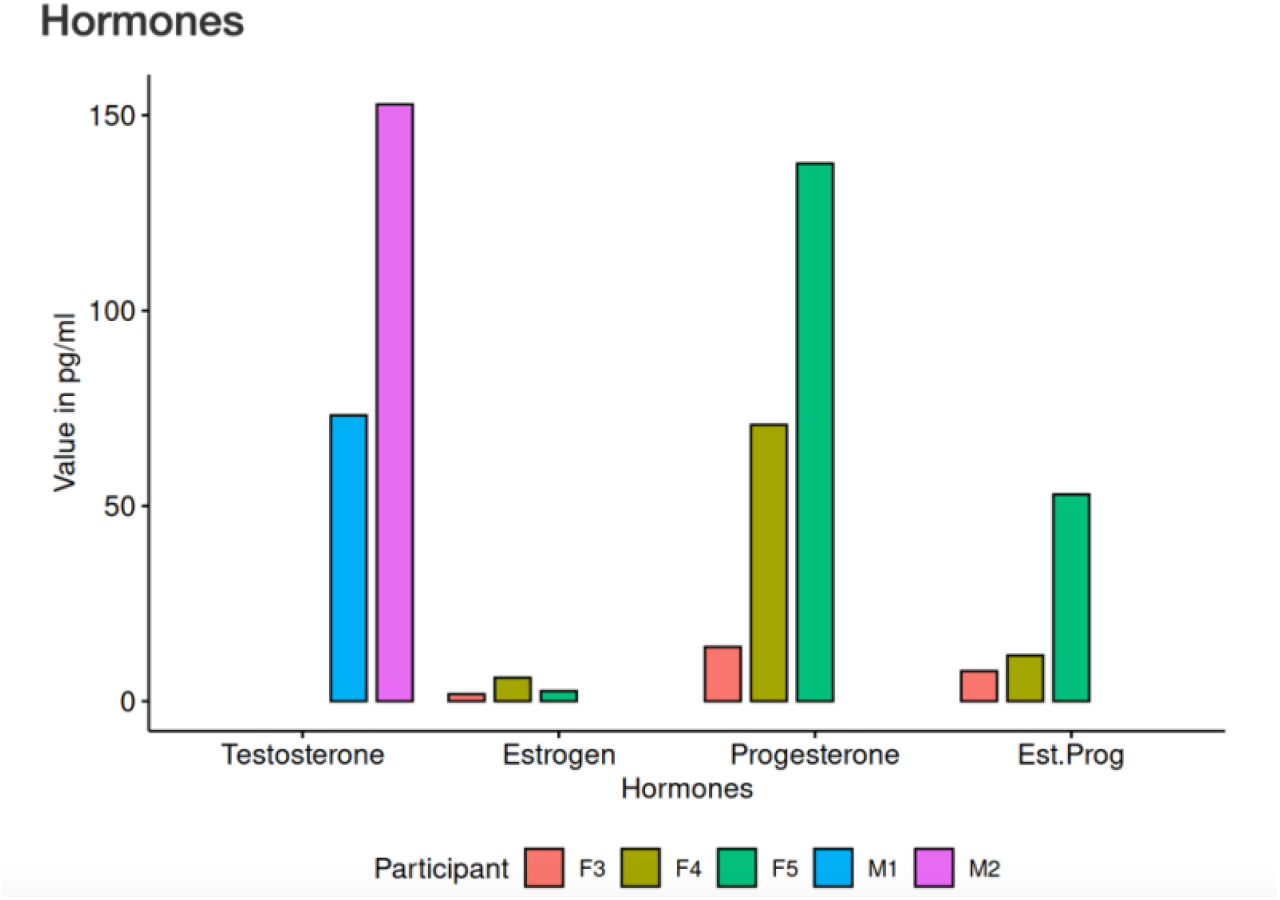
Hormonal measurement values upon wakening: testosterone for male participants and estrogen/progesterone for female participants. Females are designated with the letter ‘F’ followed by a single digit, consistent with other tables in this study, while males use the letter ‘M’ with the same numeric system.

The estrogen and progesterone levels in the female subjects varied, landing between the normal and lower end of the standard reference values, analyzed according to phases of the women’s menstrual cycles (see Table 3). In this study, the female participants were not required to take the hormonal measurement during a specific phase of their menstrual cycle, as the team wanted to make a more general analysis of estrogen and progesterone.

**Table 3.** Reference values of estrogen levels (left), progesterone (middle) and ratio progesterone / estrogen (right) in women, as referenced from laboratory values and the website GetTested.

|  |  |  |  |  |  |
| --- | --- | --- | --- | --- | --- |
| Follicular phase | 3,1-6,4 pg/ml | Follicular phase | 30,3 - 51,3 pg/ml | Follicular phase | 4-14 |
| Ovulation phase | 4,9-11,9 pg/ml | Luteal phase | 87,3 - 544,3 pg/ml | Luteal phase | 10-131 |
| Luteal phase | 3,6-7,5 pg/ml | Postmenopause | 21,0 - 69,0 pg/ml | Postmenopause | 2-20 |
| Postmenopause | 3,0-7,5 pg/ml |  |  |  |  |

As women’s hormonal health has been gaining prominence in recent research [27], there’s a heightened need for prompt and accessible hormonal testing for gonadal hormones, including estrogen and progesterone, on a global scale.

As mentioned in the preceding sections, the sensitivity of saliva as a biomarker and the potential challenges in handling and delivery could have slightly influenced the female hormonal results for one of the participants who experienced abnormalities in these biomarkers (referring to Figure 1). Without further examinations being made, it is challenging to pinpoint the exact cause of these anomalies or determine if they reflect genuine physiological variations or are simply the result of external factors related to sample collection and processing.

In the study, associated to the cognitive task, three out of the five participants (3/5) exhibited a significant increase in cortisol in the 5-hour post-wakening measurement (see Figure 2) — a spike not typically expected at that time of day [20]. This unexpected rise in cortisol levels, even in a controlled setting, suggests that everyday stressors can have tangible physiological effects on the body. A potential faulty measurement was also detected in one of the participant’s final cortisol samples (referring night measurement in Figure 2, person 3), most likely due to reasons like those mentioned in the subsequent sections discussing hormonal anomalies.

**Figure 2.**
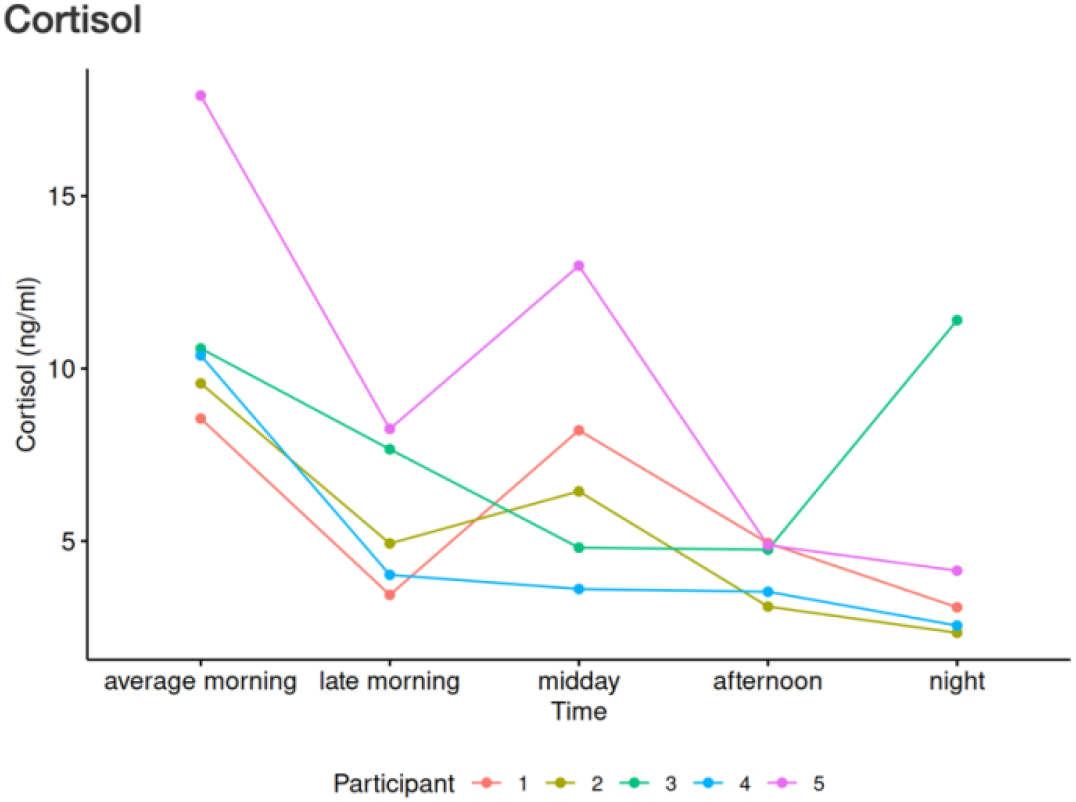
Hormonal measurement values of cortisol for all female and male participants in the study. Seven saliva samples were taken in total, with a mean value formed from the first three measurements taken during the morning hours. Given the varied waking times of the participants (ranging from 6:00-9:00 am), the data is presented in general time blocks rather than exact timestamps. Participants 1-2 = male, participants 3-5 = female.

The interesting cortisol result highlights the importance of finding effective ways to manage even minor, stressful situations in everyday scenarios. It also underscores the value of building resilience in order to conquer and thrive in daily challenges. Furthermore, it opens up avenues for further research into the links between daily stressors, cortisol response, and cognitive performance, as well as applying neurologically supported protocols to optimize individual responses to stress.

### Cardiovascular Measurements of the smart ring

By analyzing the cardiovascular measurements during the night prior to the mental task, the aim was to identify potential connections between sleeping patterns anticipating an unexpected event, referring to prior studies creating associations between changes in heart rate variability and heightened cortisol levels [28], [29]. Based on the provided data taken from the participants’ Oura application, the following results are being presented in Figure 3.

**Figure 3.**
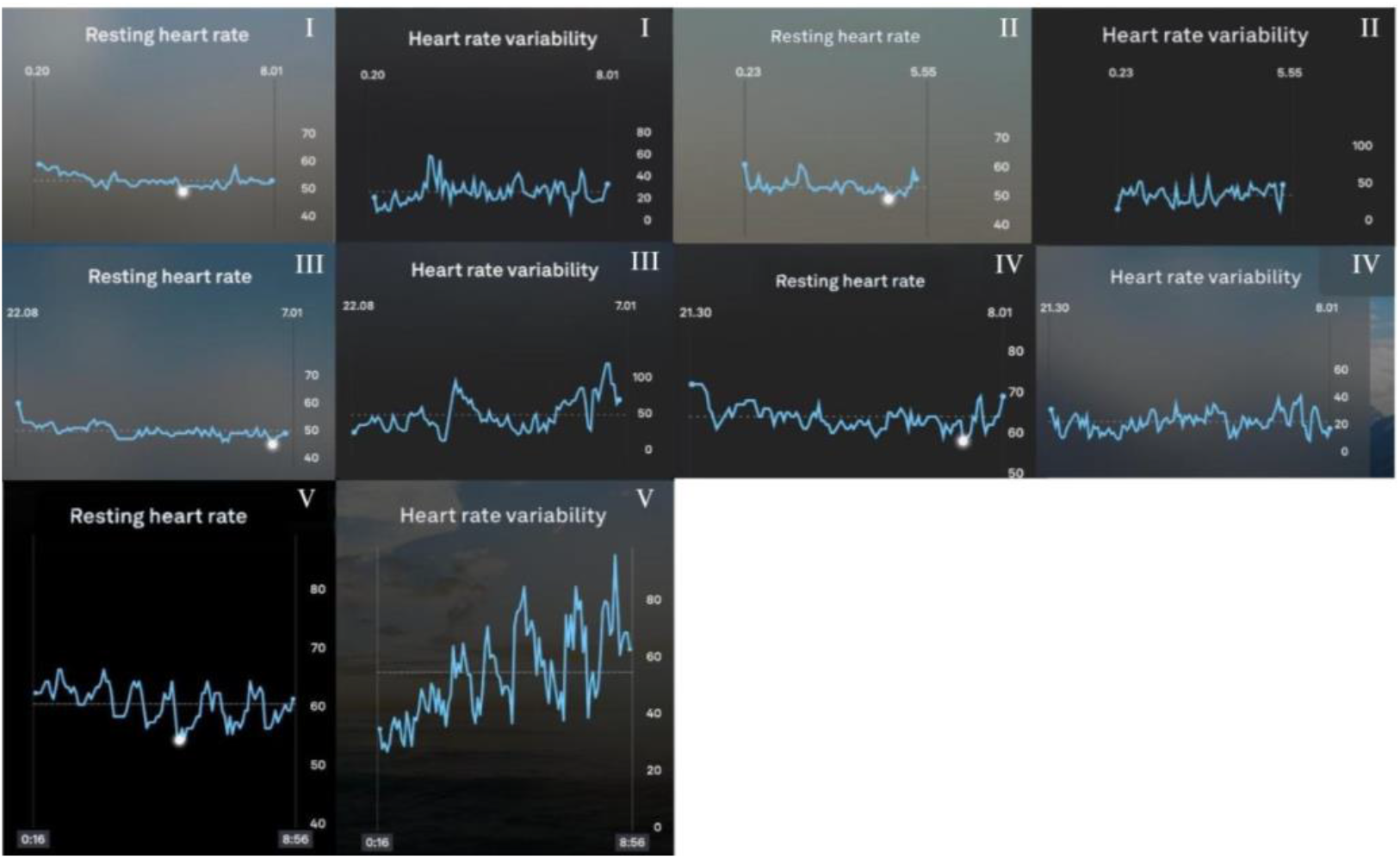
Cardiovascular rates sourced from the participants’ Oura application, highlighting resting heart rate and heart rate variability results from the night preceding the cognitive task and biometric measurements day. Participants 1-2 = male, participants 3-5 = female.

Building upon prior studies [28], [29], the study’s findings explored associations between changes in nighttime HRV and post-task cortisol levels. Those with greater changes in HRV typically demonstrated heightened cortisol responses post-task (see Figure 2), underscoring the close relationship between cardiovascular indicators of stress and the endocrine system [30]. It’s important to note, however, that these findings are derived from a relatively small set of participants. As such, while the results provide a foundational insight into the potential relationship between nighttime HRV changes and cortisol responses, they should be interpreted with caution. Further studies with larger sample sizes are recommended to validate and expand upon these preliminary findings.

### Cardiovascular Measurements of the Pulse Oximeters

Pulse oximeters, while widely used in many studies, have occasionally been noted for presenting offset results in initial measurements [31], aligning with the observations made in this study (see Figure 4). A slight drop in blood oxygen saturation was observed in three out of five (3/5) participants during the mental task. This mirrors findings from past research, suggesting that cognitive challenges can induce subtle alterations in oxygen levels [32]. Four out of five of the subjects in the study experienced an increase in their resting heart rate as a physiological response to the challenge presented. This cardiovascular result is also consistent with findings from previous studies in the field [33].

**Figure 4.**
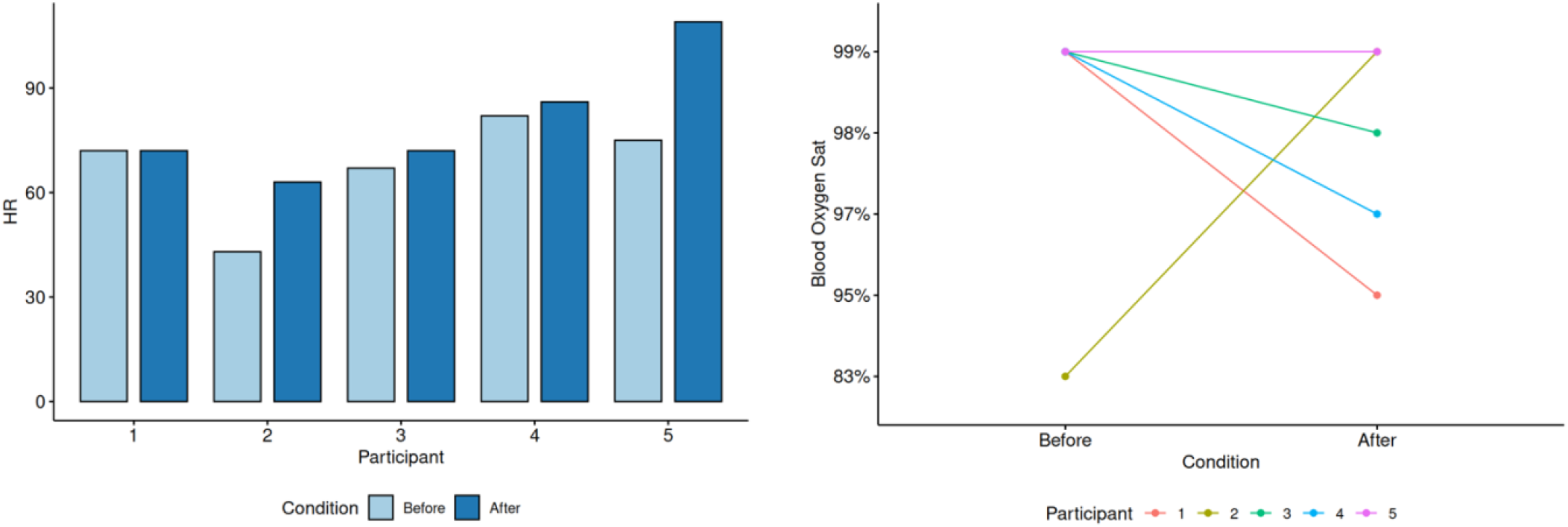
Resting heart rate measurements (left) and blood oxygen saturation levels (right) during the cognitive task from all participants. Participants 1-2 = male, participants 3-5 = female.

For one of the participants, a significant increase in blood oxygen saturation was detected after the mental task, which goes against the overall trend observed in this research trial. This discrepancy may most likely be due to a failed measurement by the pulse oximeter used, though it’s challenging to determine with certainty without doing a more comprehensive examination.

### Cognitive Assignment

Setting aside a single participant who deviated from the general trend, individuals with higher cortisol levels during the stress-induced task generally completed less tasks than those with lower cortisol levels (see Table 4 in this section). However, due to the limited sample size of this study, it is essential to consider potential confounding factors and individual variations.

**Table 4.** Numbers of tasks completed by the participants in the 5-minute cognitive task.

|  | Mathematical Problems (~1 min) | Rapid Country Naming (~20 s) | Word Scrabble (~1min) |
| --- | --- | --- | --- |
| person 1 (male) | 1 | 1 | 2 |
| person 2 (male) | 5 | 5 | 6 |
| person 3 (female) | 2 | 2 | 3 |
| person 4 (female) | 3 | 4 | 5 |
| person 5 (female) | 2 | 2 | 3 |

Therefore, suggesting the need for further longitudinal studies to determine causality and the potential long-term effects of chronic stress on cognitive performance. Individuals who reported heightened feelings of stress before the task and expressed disappointment with their performance midway through the assignments, were also the ones who generally performed at a lower level on the tasks. As demonstrated in prior research [34], [35], this indicates that individuals’ mindsets can have a powerful impact on cognitive performance. Additionally, it emphasizes the importance of self-talk in order to cultivate and maintain an optimal mindset throughout daily situations perceived as challenging. The mental task reinforces the theory that acute stress, combined with an unsettling feeling, can temporarily compromise certain cognitive functions [23], [30].

Overall, the core insights and preliminary data of biomarkers behavioral patterns found in this small-scaled study, lay the groundwork for the team’s subsequent research endeavors, to be elaborated upon in the concluding section.

## Discussion & Future advancements

With mental health issues substantially increasing after the pandemic [36], several factors have come forward as potential contributors to this alarming trend. Financial instability from economic disruptions [37], prolonged periods of isolation due to pandemic-related measures [36], and the global political tensions and uncertainties of recent years [38], serve as prime examples of factors affecting mental well-being worldwide. Yet, the paradoxical observation is that mental health-related applications and software are being produced at an exponential rate during this time [39].

Many HealthTech solutions are becoming progressively accurate when it comes to measuring and tracking health data, but in most cases they are not necessarily taking into consideration the psychological aspect associated with mental challenges. Merely analyzing the heart rate of a person who may have unhealthy, deep-rooted beliefs, might necessarily not help them to the extent that they can break those thought patterns in order to do long-term improvements in their health. Instead, it is important to consider integrating both a neuroscientific and psychological component into forthcoming solutions for a more comprehensive and multidisciplinary approach that can benefit a broader population. Having more deliberate control over one’s internal well-being and being able to manage uncomfortable or stressful situations everyone inevitably face on a daily basis, is of urgent need in our current society.

Combining the latest advancements of biosensing technologies within in a novel device with deliberate actions taken from the users themselves, is what the research team believes is the key to bridging the gap between mental health challenges and the potential solutions that the innovative field of HealthTech can offer.

The slightly unstable results from the pulse oximeters for one of the male subjects, as well as in one of the female hormonal tests (see Figures 1 and 4), emphasizes the need of developing alternative, new solutions to conveniently and with a higher accuracy measure cardiovascular and hormonal data in real-time. From the personal testimonials of the research study’s participants, several challenges were highlighted. They expressed the inconvenience of sequentially providing saliva samples and sending them for laboratory analysis. Additionally, enduring long wait times and trusting the accuracy of the results were concerns that also were mentioned. The minor complications of getting an initial value measured from the pulse oximeter devices were also addressed. Based on these insights, this preliminary study suggests a pressing need for innovation. Modern technology and machine learning could streamline the process, making the detection and prevention of unfavorable physiological conditions easier. Consequently, the study also raises the question of whether the subjects in this initial research trial might have improved their performance on the mental task assignments if they had been introduced to neurologically validated techniques to alter their physiological state [40], [41].

Developing a device intended to be used as a tool to have at hand when feelings of discomfort arise or mental focus is needed instead of having to wear it continuously, is something that could be of high value due to the increased flexibility and freedom. We are approaching a juncture where such a device might soon be seeing the light of day, offering well-rounded insights and guidance to our physiological and psychological well-being, and innovating remote healthcare as we know it.

With this initial research study being completed and getting a comprehensive understanding of biomarkers behavior and interference with cognitive function, the research team will continue to delve deeper into the details of these interactions, aiming to refine methodologies and expand the participant group to further validate and generalize the findings to a broader population.

## Limitations of the Study

While this study provides insights into the physiological patterns under specific conditions, several limitations should be acknowledged. Firstly, as this was a pilot study aimed at exploring theories and gaining knowledge, the sample size was notably small, comprising only five participants.

Although this facilitated an individualized analysis, the limited sample size restricts the external validity and broader applicability of the findings. It is, therefore, imperative that results are interpreted with caution when considering their relevance to a more extensive population.

Secondly, certain cardiovascular and hormonal measurements presented minor inconsistencies. These irregularities could arise from instrumental variances or slight differences in procedural execution. As such, the potential margin of error must be considered when assessing the precision and reliability of the results. To reduce these inconsistencies, future studies should adopt strict calibration and standardization protocols and be conducted on a larger scale.

## Rationale for Utilizing Oura Data

In the research study, the team opted to utilize data from the Oura Ring, a well-established tool in personal health tracking. Oura permits users to export comprehensive data in formats such as CSV, JSON, and summarized PDF reports. Specifically, the ‘Share Report’ function provides a detailed summary of sleep trends, including sleep stages, total sleep time, wake-up times, and daily movement, available in timeframes of seven, 30, or 90 days. The ability to retrieve this data in standardized formats ensures consistency in the trial’s analysis. Given the accessibility and granularity of the data available through Oura, its inclusion in this study serves to enhance the robustness of the consequent findings.

## Acknowledgments

We gratefully acknowledge the participants for generously volunteering their time and effort, and we deeply appreciate the great collaboration with the study’s data analyst, Rishi Das Roy.

